# Development and validation of a triplex qPCR assay to detect efflux pump-mediated antibiotic resistance in *Burkholderia pseudomallei*

**DOI:** 10.1101/301960

**Authors:** Jessica R. Webb, Erin P. Price, Nawarat Somprasong, Herbert P. Schweizer, Robert W. Baird, Bart J. Currie, Derek S. Sarovich

**Affiliations:** Global and Tropical Health Division, Menzies School of Health Research, Darwin, Northern Territory, Australia; Faculty of Science, Health, Education and Engineering, University of the Sunshine Coast, Sippy Downs, Queensland, Australia; College of Medicine, Department of Molecular Genetics and Microbiology, Emerging Pathogens Institute, University of Florida, Gainesville, Florida, USA; Departments of Infectious Diseases and Pathology and Northern Territory Medical Program, Royal Darwin Hospital, Darwin, Northern Territory, Australia

**Keywords:** *Burkholderiapseudomallei*, antibiotic resistance, melioidosis, efflux pump, gene expression, qPCR

## Abstract

*Burkholderia pseudomallei*, the causative agent of the deadly tropical disease melioidosis, is intrinsically resistant to many antibiotics, leaving few effective treatment options. Trimethoprim-sulfamethoxazole (SXT), meropenem (MEM) and doxycycline (DOX) are valuable antibiotics for melioidosis treatment due to inherently low or no primary resistance. Although considered rare, upregulation of one or more resistance-nodulation-division (RND) efflux pumps is now known to lead to acquired resistance towards these drugs in *B. pseudomallei.* Here, we developed a triplex quantitative PCR assay to detect upregulation of the three clinically relevant RND efflux systems: AmrAB-OprA, BpeB-OprB and BpeEF-OprC. The triplex assay was tested on seven clinically-derived *B. pseudomallei* isogenic pairs, where the latter strain of each pair had altered regulator activity and exhibited reduced susceptibility to SXT, MEM or DOX. The triplex assay accurately detected efflux pump upregulation between isogenic pairs, which corresponded with decreased antibiotic susceptibility. We further verified assay performance on eight laboratory-generated *B. pseudomallei* mutants encoding efflux pump regulator mutations. Targeting antibiotic resistance in *B. pseudomallei* using molecular genotyping provides clinicians with a rapid tool to identify potential treatment failure in near real-time, enabling informed alteration of treatment during an infection and improved patient outcomes.

**IMPORTANCE:** The melioidosis bacterium *Burkholderia pseudomallei* is intrinsically resistant to many antibiotics, limiting treatment options to a handful of drugs including meropenem, doxycycline and trimethoprim-sulfamethoxazole. Although rare, there have now been several documented melioidosis cases where resistance to these antibiotics has developed during an infection, leading to treatment failure and increased mortality rates. Interestingly, all strains resistant to these drugs exhibit increased efflux pump expression, representing a shared molecular signature that can be exploited for rapid diagnostic purposes. Here, we developed and validated a single-tube real-time qPCR assay to detect clinically relevant efflux pump upregulation in *B. pseudomallei*, an important first step towards high-level resistance. This triplex assay offers a drastically reduced turn-around-time compared to current methodology, enabling earlier detection of resistance emergence. Implementation of this new diagnostic will aid clinicians in the selection of appropriate therapy, thereby minimizing resistance development and treatment failure for this high-mortality disease.

## INTRODUCTION

The development of antibiotic resistance in Gram-negative bacteria has become a global crisis as declared by the World Health Organization (1) and Centers for Disease Control and Prevention (2). The Gram-negative bacterium *B. pseudomallei*, the causative agent of melioidosis, is one of the most intrinsically antibiotic resistant bacteria (3, 4, 5), due to its diverse array of chromosomally-encoded resistance mechanisms (6). This inherent resistance limits melioidosis treatment options to a small number of antibiotics. Resistance emergence to any of these drugs during treatment is thus of great concern given the limited number of alternative treatments.

Melioidosis is arguably one of the most neglected tropical diseases of our time, with gross underreporting of cases, particularly in emerging endemic regions (7). Recent modelling suggests that there are 165,000 melioidosis cases worldwide annually, of which 89,000 are fatal, rates that are similar to the much higher profile disease, measles (8). There is currently no vaccine towards *B. pseudomallei*, with treatment fully reliant on antibiotic administration, which is biphasic and lengthy. Treatment typically involves 10-14 days of intravenous ceftazidime (CAZ), with meropenem (MEM) used in severe cases or when resistance develops towards CAZ (9). The eradication phase consists of at least three months of trimethoprim-sulfamethoxazole (SXT) or amoxicillin-clavulanate, with doxycycline (DOX) used in instances where patients develop impaired renal function, bone marrow suppression or skin reactions due to SXT intolerance (10). The emergence of resistance during treatment is fortunately rare, but has now been documented for almost all antibiotics used to treat melioidosis (11,12,13,14,15), the exception being high-level (>12 μg/mL) MEM resistance. Alarmingly, cases of *B. pseudomallei* exhibiting decreased susceptibility to MEM have recently been identified in Australian patients (16), and are associated with prolonged blood culture positivity and poorer patient outcomes (17).

*B. pseudomallei* encodes for three clinically relevant resistance-nodulation-cell division (RND) efflux pumps: AmrAB-OprA (*BSPL1802*-*BPSL1804*), BpeAB-OprB (*BPSL0814*-*BPSL0816*) and BpeEF-OprC (*BPSS0292*-*BPSS0294*), which give rise to resistance towards multiple antibiotic classes (18,19,20). Each RND system consists of a membrane fusion protein (AmrA, BpeA, and BpeE, respectively), an RND transporter (AmrB, BpeB, and BpeF, respectively), an outer membrane protein (OprA, OprB, and OprC, respectively), and one or more regulators (AmrR [BPSL1805], BpeR [BPSL0812], and BpeT [BPSS0290] and BpeS [BPSL0731], respectively) (21). Overexpression of these efflux pumps can occur during the course of melioidosis treatment due to loss-of-function mutations in repressors or mutations leading to co-inducer independence of activators in their associated regulatory genes (17, 22, 23, 24). For instance, certain mutations affecting theTetR-type regulator gene, *amrR*, cause decreased MEM susceptibilities in melioidosis patients with prolonged infections (17), and in combination with mutations in the SAM-dependent methyltransferase gene *BPSL3085*, can lead to clinically significant DOX resistance (23). The *bpeT* and *bpeS* genes encode two closely related LysR-type transcriptional regulators. Mutations that affect the carboxy-terminal co-inducer-binding domains of BpeT and BpeS, usually together with mutations in a tetrahydrofolate pathway-linked pterin reductase gene, *ptr1/folM*, have been associated with SXT resistance (22, 24). The *bpeR* regulator is less well-characterized, although a mutation within this gene has been found in a clinical *B. pseudomallei* isolate that has intermediate resistance to multiple drugs, including MEM (17). Importantly, all observed cases of MEM, DOX and SXT resistance have involved upregulation of one or more of these three RND efflux pumps (17, 25).

Despite the critical role that these RND efflux systems play in conferring acquired antibiotic resistance in *B. pseudomallei*, no methods are currently available to quickly and simultaneously detect their altered expression. A number of singleplex PCR assays have been published (24, 26, 27, 28); however, these assays are SYBR Green-based and thus cannot be multiplexed, making them more time-consuming and costly to perform. Additionally, SYBR Green-based assays are typically less robust and less specific than probe-based PCR assays due to the non-specific nature of the fluorogenic SYBR Green dye, which detects all double-stranded DNA molecules. Therefore, the aim of this study was to develop a robust and highly specific fluorogenic probe-based triplex quantitative real-time PCR(qPCR) assay to simultaneously detect the upregulation of the RND efflux pumps AmrAB-OprA, BpeAB-OprB and BpeEF-OprC in strains exhibiting increased minimum inhibitory concentrations (MICs) towards DOX, MEM and SXT. The triplex assay was first validated to determine the limits of quantitation (LoQ), detection (LoD), and linearity, and then tested against a panel of genetically characterized, antibiotic resistant *B. pseudomallei* isolates.

## RESULTS

### Limits of quantitation (LoQ) and detection (LoD), and linearity of the triplex qPCR assay

Following the design of probe-based assays targeting *amrB*, *bpeB* and *bpeF*, the lower LoQ and LoD was first calculated for each assay in the singleplex format, and subsequently in the triplex format. LoD was defined as the lowest analyte concentration at which detection is feasible, and LoQ was defined as the lowest concentration of analyte that can be determined with an acceptable level of precision and accuracy (29). Based on these definitions, the LoQ was determined as the lowest amount of DNA where 8/8 replicates amplified with a CT standard deviation (*σ*) of <0.8, and with good efficiency (R^2^ values >0.98), and LoD was defined as the concentration where at least 2/8 replicates amplified, irrespective of *σ* or efficiency. The lower LoQ in the singleplex format for all three assays was ≥4×10^‒4^ ng (≥400 fg, 52 genomic equivalents (GEs)), and the LoD was ≥4×10^‒6^ ng (≥4 fg, 0.5 GEs), ≥4×10^‒5^ ng (≥40 fg, 5 GEs) and ≥4×10^‒5^ ng (≥40 fg, 5 GEs) for *amrB*, *bpeF* and *bpeB*, respectively. In the triplex format, for all targets, the LoQ was ≥4×10^‒3^ ng (≥4 pg, 515 GEs), and the LoD was ≥4×10^‒5^ ng (≥40 fg, 5 GEs) (Figure 1).

**FIG 1.**
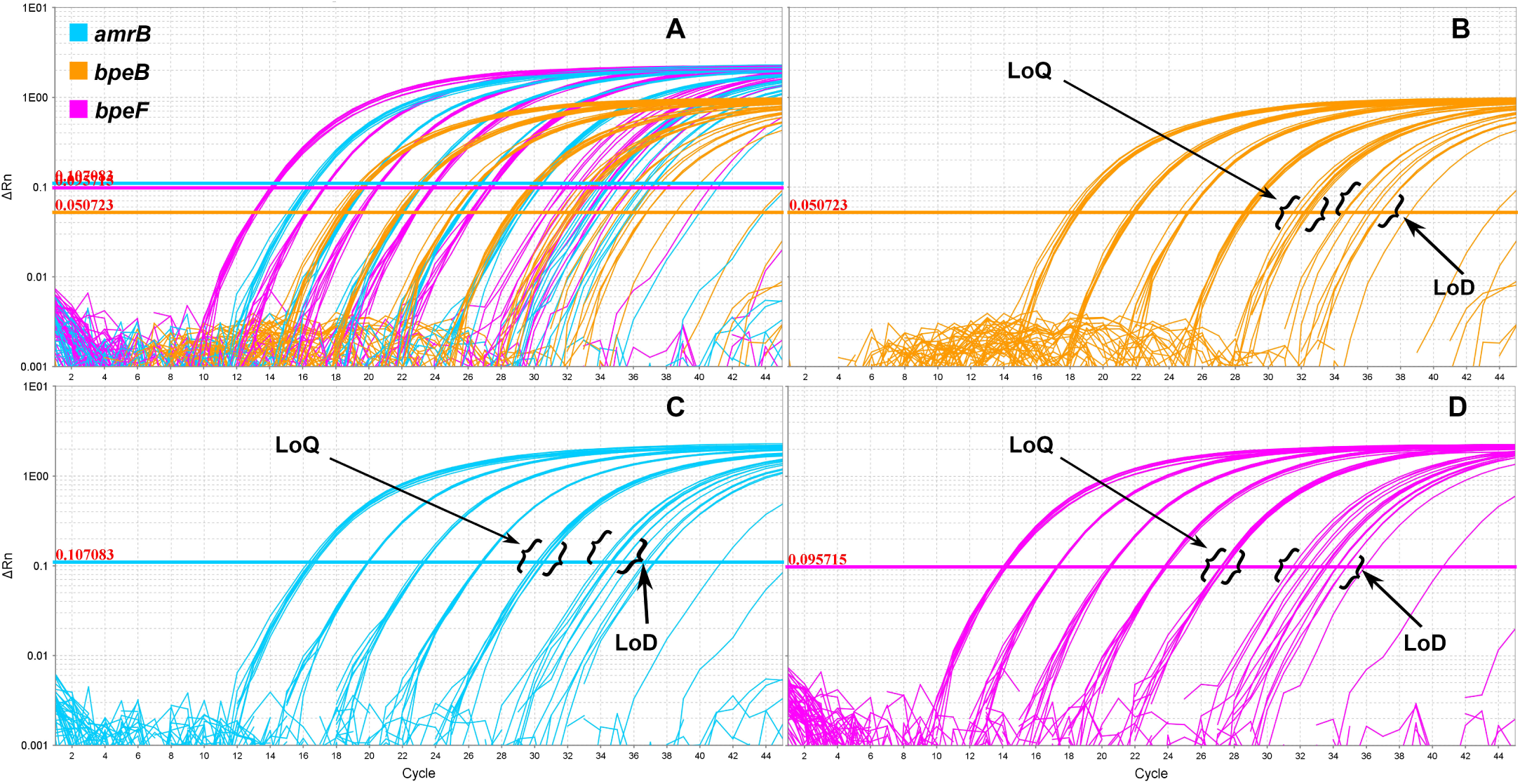
Quantitative PCR amplification plots of the *Burkholderia pseudomallei* resistance-nodulation-division efflux pump triplex assay. A) Amplification across eight replicates at DNA concentrations ranging from 40 to 4×10-6 ng. B) Amplification plot of *bpeB.* C) Amplification plot of *amrB.* D) Amplification plot of *bpeF.* LoQ, limit of quantitation;LoD, limit of detection.

Next, the linearity of these assays was determined in the triplex and singleplex format. Efficiency (linearity) was measured by the rate at which a PCR amplicon is generated (30), where the maximum quantitative accuracy occurs in assays that function at or near 100% efficiency. In the triplex format, the standard curves for *amrB*, *bpeF* and *bpeB* all had an R^2^ value of >0.95 and showed good amplification efficiencies (90% for all three targets) (Figure S1). In singleplex format, the R^2^ values were >0.98, with an efficiency of >98% for all three assays (Figure S2).

### The conserved genes *mmsA* and 23S rDNA are suitable housekeeping genes for normalized expression analysis in *B. pseudomallei.*

To determine the performance of the conserved gene *mmsA* as an expression normalization control for all clinical isogenic and Bp82 pairs, we compared normalized efflux pump expression against both 23S and *mmsA* in the eight Bp82 mutants. The normalized fold change of the triplex efflux assay for the eight Bp82 pairs was consistent when either 23S or *mmsA* was used for normalization (Table S1), suggesting that *mmsA* expression, at least across the conditions that were tested in this study, is uniform. These results support the use of *mmsA* as a single-copy normalization control gene in *B. pseudomallei.*

### Comparative genomic analysis identifies a novel *amrR* mutation in the latter P1048 isolate

SPANDx analysis of MSHR9766 and MSHR9872 was carried out to identify the genetic basis for decreased MEM susceptibility in the latter isolate. Two missense mutations were detected in MSHR9872: a mutation in AmrR (AmrR_G50E_), and a mutation in the isoleucine tRNA synthetase gene, *ileS* (*BPSL0906*; NeS_H505G_), the product of which catalyzes the aminoacylation of Ile-tRNA. In addition, we detected a frameshift mutation in MSHR9872 within *BPSS2161* (*BPSS2161*_A28fs_), which encodes for a propanoate metabolism protein belonging to the MmgE-PrpD family. No other mutations (i.e. gene acquisition or loss, gene copy number variation) were identified between this pair.

### Efflux pump upregulation in eight clinical strains with regulatory mutations

The triplex qPCR assay was tested on eight clinical *B. pseudomallei* isolates that encompassed *amrR*, *bpeR* or *bpeT* mutants, including MSHR0052 AmrR_E190_^⋆^, which lacks an isogenic pair. None of the clinical strains tested in our study encoded *bpeS* mutations. The triplex assay showed increased efflux expression in at least one of the RND efflux pumps in those isolates containing *amrR*, *bpeR* or *bpeT* regulatory mutations (Figure 2), consistent with their role in locally coordinating efflux pump expression. The latter strain from P215, MSHR0937, which encodes a mutation within the *bpeAB*-*oprB* regulator, *bpeR* (BpeR_D176A_), had a corresponding increase in the expression of *bpeB* (15x;3.8-fold;Figure 2A) compared with its WT isogenic pair, MSHR0664. Similarly, the latter strain isolated from Patient CF6, MSHR5654, which encodes a mutation within the *bpeEF*-*oprC* regulator, *bpeT* (BpeT_T314fs_), showed significant upregulation of *bpeF* (9.5x;3.2-fold) when compared with its WT isogenic pair, MSHR5651 (Figure 2B).

**FIG 2.**
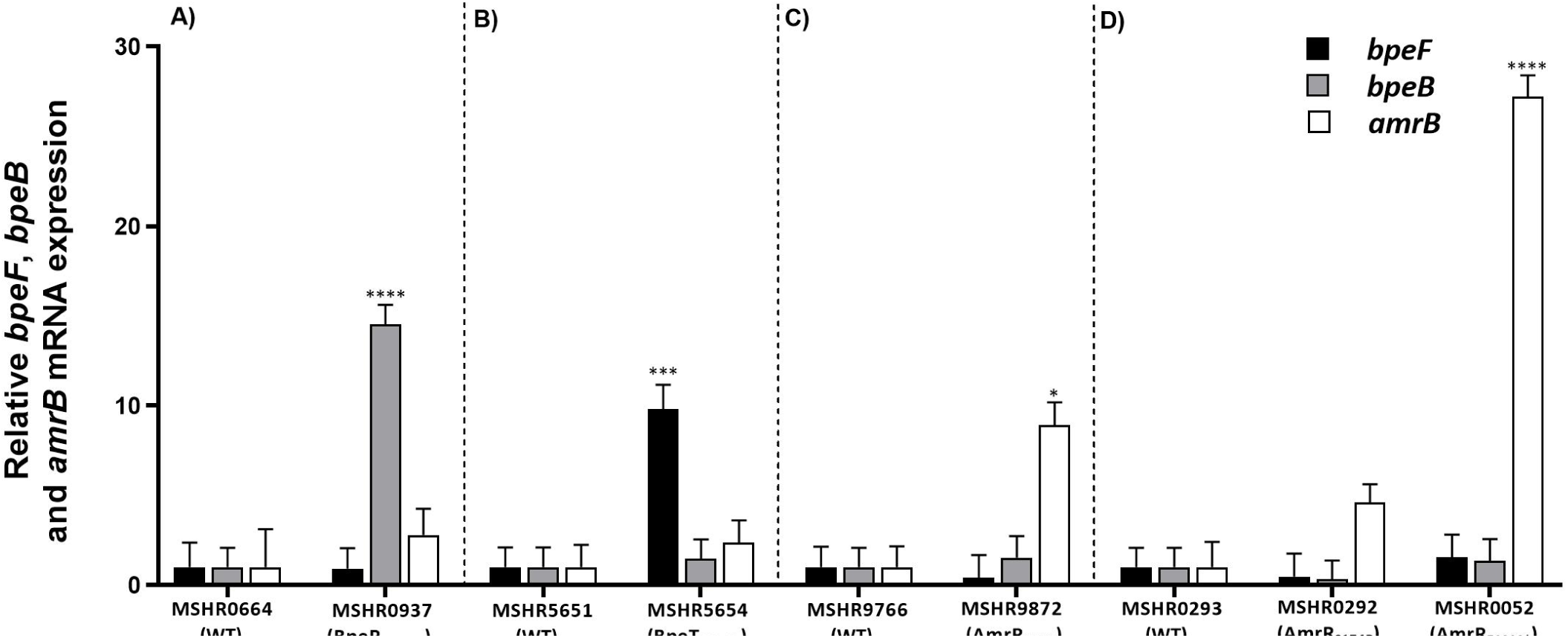
AmrAB-OprA (*amrB*), BpeAB-OprB (*bpeB*) and BpeEF-OprC (*bpeF*) expression in five melioidosis cases harboring *B. pseudomallei* isolates that encode efflux pump regulator mutations. As an isogenic WT *B. pseudomallei* pair is lacking for MSHR0052, MSHR0293 was used as the WT control for normalized efflux expression of this strain. For all other strains, the WT isogenic pair was used for normalization. Efflux pump expression was normalized against the conserved gene, *mmsA* (31). Error bars denote standard deviation between biological replicates, which were all performed in technical duplicates. The y axis represents the relative times (x) change in *amrB*, *bpeB* and *bpeF* expression. Statistical analysis was done by two-way ANOVA and Tukey’s multiple-comparison test. ^⋆⋆⋆⋆^, *p* < 0.0001; ^⋆⋆⋆^, *p* < 0.001; ^⋆⋆^ *p* < 0.01; ^⋆^, *p* < 0.05.

Six latter strains, all of which exhibited decreased susceptibility towards MEM (3-8 μg/mL; Table 1), encoded mutations within the *amrAB*-*oprA* regulator, *amrR.* When these six strains were tested with the triplex qPCR assay, only three (MSHR9872 AmrR_G50E_, MSHR0292 AmrR_S174p_ and MSHR0052 AmrR_E190_^⋆^) showed a significant increase in *amrB* expression, which ranged from 5x (2.3-fold; MSHR0292 AmrR_S174p_) to 27x (4.7-fold; MSHR0052 AmrR_E190_^⋆^) (Figures 2C and 2D). The remaining three *amrR* mutant strains, MSHR7929 AmrR_E30D_, MSHR4083 AmrR_Δ_A153-D156__ and MSHR6755 AmrR_Δ_V60-C63__, showed no significant increase in *amrB* expression. Importantly, the amplification of the triplex assay for all the isogenic strains was within the LoQ of the triplex assay, ruling this factor out as a cause for the lack of differential expression.

**TABLE 1.** *Burkholderiapseudomallei* clinical isolates and laboratory-generated mutants used in this study. Mutations in resistance-nodulation-division efflux regulatory genes are indicated. Minimum inhibitory concentrations (MICs; μg/mL) for wild-type and mutant strains are shown for each antibiotic. MICs above the intermediate (i) or resistant (r) threshold determined for *B. pseudomallei* are indicated.

| Patient | Strain | Regulatory mutation | DOX | TMP | SMX | SXT | CAZ | MEM<sup>a</sup> | Study |
| --- | --- | --- | --- | --- | --- | --- | --- | --- | --- |
| **Clinical strains** |  |  |  |  |  |  |  |  |  |
| Pre-DPMS 89 | MSHR0052 | AmrR<sub>E190</sub><sup>b</sup> | 48 (r) | ND | ND | ND | ND | 8 (i) | (17) |
| Non-DPMS 00 | MSHR0293 | WT | 1 | ND | ND | 0.38 | 1 | 0.5 | (23) |
|  | MSHR0292 | AmrR<sub>S174P</sub> | 16 (r) | ND | ND | 0.50 | 1 | 2 |  |
| CF6 | MSHR5651 | WT | 0.38 | ND | ND | 0.75 | 1.5 | 0.5 | (22) |
|  | MSHR5654 | BpeT<sub>T314fs</sub> | 3 | ND | ND | >32 (r) | >256 (r) | 2 |  |
| P215 | MSHR0664 | WT | ND | ND | ND | 2 | 0.75 | 1.5 | (17) |
|  | MSHR0937 | BpeR<sub>N176A</sub> | 4 | ND | ND | 2 | 8 | 6 (i) |  |
| P1048 | MSHR9766 | WT | ND | ND | ND | ND | ND | 0.5 | This study |
|  | MSHR9872 | AmrR<sub>G50E</sub> | ND | ND | ND | ND | ND | 6 (i) |  |
| P797 | MSHR6522 | WT | 1 | ND | ND | 1.5 | 1.5 | 0.5 | (17) |
|  | MSHR7929 | AmrR<sub>E30D</sub> | 2 | ND | ND | 4 (r) | 1.5 | 4 |  |
| P608 | MSHR3763 | WT | 0.75 | ND | ND | 3 (i) | 2 | 0.75 | (17) |
|  | MSHR4083 | AmrR<sub>ΔA153-D156</sub> | 1 | ND | ND | 24 (r) | 2 | 6 (i) |  |
| P726 | MSHR5864 | WT | 1 | ND | ND | 1.5 | 1.5 | 0.75 | (17) |
|  | MSHR6755 | AmrR<sub>ΔV60-C63</sub> | 1.5 | ND | ND | 0.75 | 1.5 | 3 |  |
|  | MSHR6755 | Δ*amrR* | ND | ND | ND | ND | ND | 3 |  |
| **Bp82 mutants** |  |  |  |  |  |  |  |  |  |
| N/A | Bp82 | WT | 0.5 | 0.75 | 4 | 0.094 | 2 | ND | (50) |
| N/A | Bp82.268 | BpeT<sub>C310R</sub> | 2 | 4 | 8 | 0.38 | 2 | ND | (24) |
| N/A | Bp82.269 | BpeT<sub>L265R</sub> | 1 | 4 | 8 | 0.38 | 2 | ND | (24) |
| N/A | Bp82.270 | BpeT<sub>S280P</sub> | 8 | >32 | ND | ND | 2 | ND | (24) |
| N/A | Bp82.284 | BpeS<sub>P29S</sub> | 2 | >32 | ND | ND | 2-4 | ND | (24) |
| N/A | Bp82.285 | BpeS<sub>K267T</sub> | 8 | >32 | ND | ND | 1-2 | ND | (24) |
| N/A | Bp82.280 | Δ*AmrAB-OprA* Δ*BpeR* | 1 | 0.75 | ND | ND | 2 | ND | (24, 27) |
| N/A | Bp82.410 | Δ*AmrR* | 4 | 0.75 | ND | ND | 2 | ND | This study |
| N/A | Bp82.412 | AmrR<sub>S174P</sub> | 4 | 0.75 | ND | ND | 2 | ND | This study |

Abbreviations: r, resistant; i, intermediate resistance; fs, frameshift mutation; ND, not determined; WT, wild type; DOX, doxycycline; TMP, trimethoprim; SMX, sulfamethoxazole; SXT, trimethoprim-sulfamethoxazole; CAZ, ceftazidime; MEM, meropenem.
<sup>a</sup>Clinical & Laboratory Standards Institute (CLSI) breakpoints for MEM have yet to be determined. Therefore, cut-offs are instead included for the related carbapenem antibiotic, imipenem.
<sup>b</sup>nonsense mutation leading to a premature stop codon.

### Induction of AmrAB-OprA in *B. pseudomallei amrR* mutants exhibiting decreased MEM susceptibilities

To better understand the lack of differential expression of *amrAB-oprA* in some strains with altered *amrR*, the triplex qPCR assay was tested on RNA extracted from MSHR4083, MSHR6755 and MSHR6755 Δ*amrR* grown in the presence of a sub-inhibitory (0.25 μg/mL) concentration of MEM. For MSHR6755 AmrR_Δ_V60–C63__, *amrB* was dramatically upregulated (24x; 4.6-fold) in the presence of MEM (Figure 3B). Additionally, the triplex assay revealed a subtle (2x; 0.9-fold) increase in the expression of *bpeB* and *bpeF* in this strain when grown in the presence of MEM (Figure 3B). In MSHR6755 Δ*amrR*, MEM induction also resulted in 12x (3.5-fold) *amrB* upregulation (Figure 3B). Similarly, MEM induction led to 21x (4.2-fold) upregulation of *amrB* in MSHR4083 AmrR_Δ_A153–D156__ (Figure 3C). Unlike MSHR6755 AmrR_Δ_V60–C63__′, MEM did not induce *bpeB* and *bpeF* upregulation in MSHR6755 Δ*amrR* or MSHR4083 AmrR_Δ_A153–D156__.

**FIG 3.**
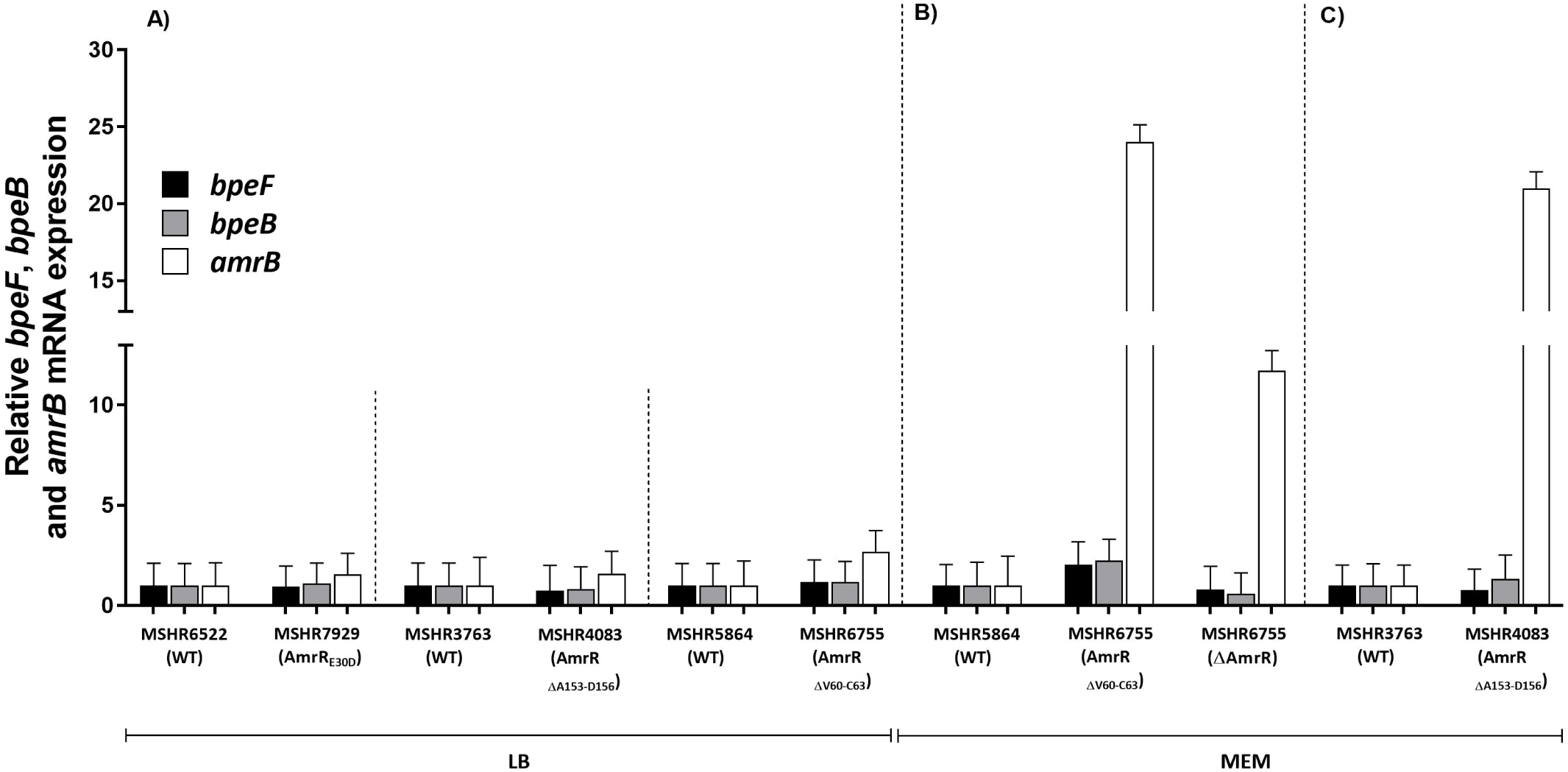
AmrAB-OprA (*amrB*), BpeAB-OprB (*bpeB*) and BpeEF-OprC (*bpeF*) expression in *Burkholderia pseudomallei*, with the latter isolate of each pair containing *amrR* point mutations or deletions that confer decreased meropenem (MEM) susceptibility. A) Three isogenic pairs grown in Luria-Bertani (LB) broth; B) isogenic pair MSHR5864 and MSHR6755 (and AmrR knockout MSHR6755 Δ*amrR*) grown in LB plus MEM; C) isogenic pair MSHR3763 and MSHR4083 grown in LB broth plus MEM. Efflux pump expression was normalized against the conserved gene, *mmsA* (31). Error bars denote standard deviation between biological duplicates, which were all performed in technical duplicates. The y axis represents the relative times (x) change in *amrB*, *bpeB* and *bpeF* expression. Statistical analysis was done by two-way ANOVA and Tukey’s multiple-comparison test, a statistically significant *p* value was not identified between pairs.

### Increased efflux expression in laboratory-generated efflux regulator mutants

The triplex qPCR assay was tested on eight laboratory-generated Bp82 mutants con-taining mutations affecting the *bpeT*, *bpeS* or *amrR* efflux pump regulators (Figure 4). The three *bpeT* missense mutants, Bp82.268 BpeT_C310R_, Bp82.269 BpeT_L265R_ and Bp82.270 BpeT_S280P_, exhibited *bpeF* upregulation of 28x (4.8-fold), 38x (5.2-fold) and 73x (6.1-fold), respectively, but no significant upregulation of the other two efflux pumps. The two *bpeS* missense mutants, Bp82.284 BpeS_P29S_ and Bp82.285 BpeS_K267T_, exhibited upregulation of *bpeF* of 89x (6.5-fold) and 145x (7.2-fold), respectively. In Bp82.284 BpeS_P29S_, *bpeB* was also moderately upregulated (~3x; 1.5-fold). Upregulation of *amrB* was confirmed in the two *amrR* mutants Bp82.412 AmrR_S174P_ (6x; 2.6-fold) and Bp82.410 Δ*amrR* (7x; 2.8-fold), with no demonstrable differential expression of *bpeB* and *bpeF.* Lastly, in Bp82.280 Δ(*amrAB*-*oprA*) Δ*bpeR*, the triplex qPCR assay showed no *amrB* amplification due to deletion of this locus, but 5x (2.3-fold) upregulation of *bpeB* due to deletion of *bpeR.* In all instances where amplicons were produced, triplex assay amplification for the isogenic Bp82 mutants was within the LoQ of the triplex assay.

**FIG 4.**
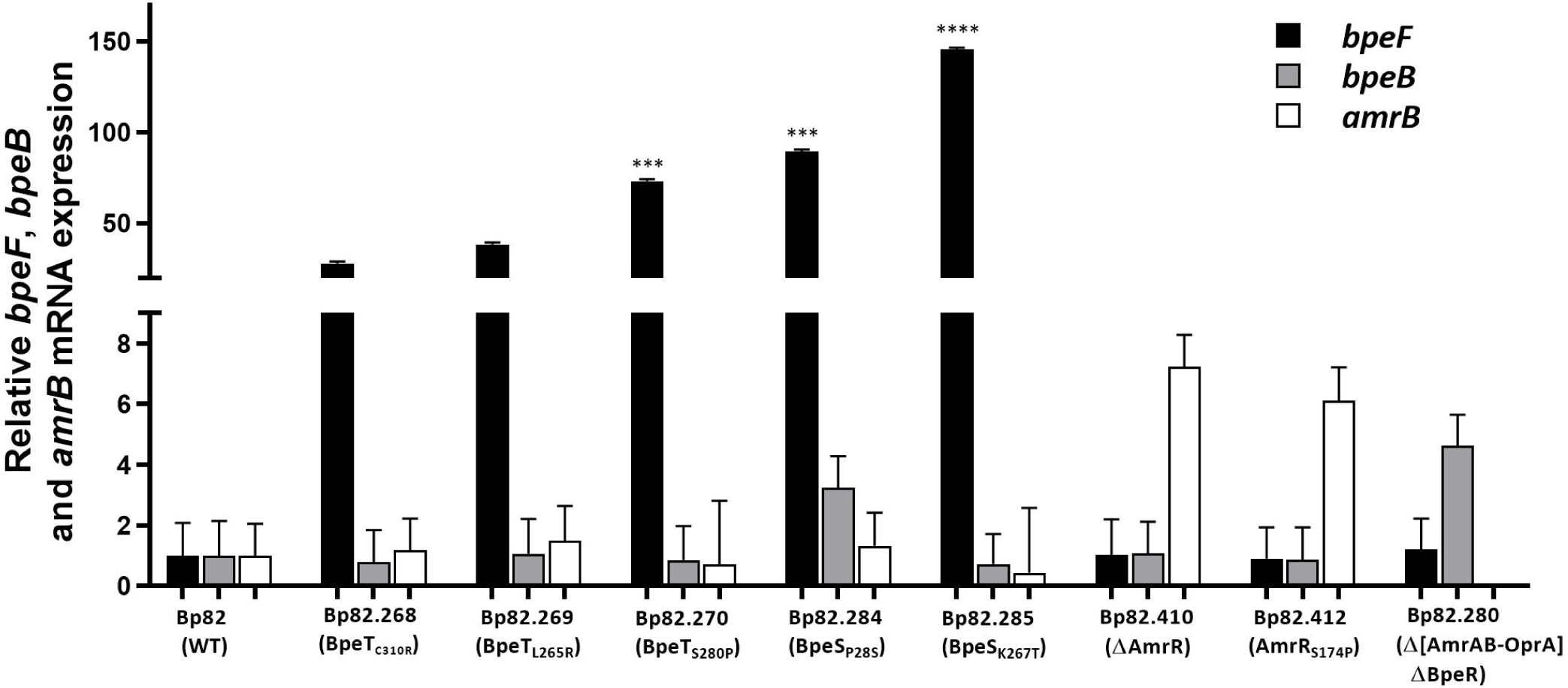
AmrAB-OprA (*amrB*), BpeAB-OprB (*bpeB*) and BpeEF-OprC (*bpeF*) expression in eight laboratory-generated *Burkholderia pseudomallei* Bp82-derived mutants containing regulator mutations relative to WT Bp82. Error bars denote standard deviation between biological triplicates, which were all performed in technical triplicates. Efflux pump expression was normalized against the conserved gene, *mmsA* (43). The y axis represents the relative times (x) change in *amrB*, *bpeB* and *bpeF* expression. Statistical analysis was done by two-way ANOVA and Tukey’s multiple-comparison test. ^⋆⋆⋆⋆^, *p* < 0.0001; ^⋆⋆⋆^, *p* < 0.001.

## DISCUSSION

Despite dramatic improvements in survival rates over the past 30 years, melioidosis continues to have a stubbornly high fatality rate, with between 10 and 40% of treated cases in the hyperendemic regions of northern Australia and Northeast Thailand, respectively, resulting in death (32). It is becoming clearer that at least some of these cases can be linked to the development of previously unrecognized resistance or decreased susceptibility towards clinically relevant antibiotics (17, 22, 33, 34). Of most concern, antibiotic resistance in *B. pseudomallei* has now been documented towards almost all antibiotics (17). Identification of an increasing number of novel acquired antibiotic resistance mechanisms in clinical *B. pseudomallei* strains, together with the broadening global presence of melioidosis-endemic regions, has increased the urgency for methods that can provide rapid detection of emerging antibiotic resistance.

In *B. pseudomallei*, antibiotic resistance determination currently relies on culture-based diagnostics including inhibition zone measurement (e.g. Etests) or serial dilutions of the antibiotic in culture media (e.g. broth or agar microdilutions) (35). Despite culture-based methods being accurate, they have lengthy turn-around-times, and do not determine the genetic basis of resistance, both of which are important considerations when determining optimal therapeutic strategies. Nucleic-acid based detection methods such as real-time PCR have the potential to circumvent many shortcomings of traditional culture-based methods, offering more rapid turn-around times and determination of the underlying resistance mechanism, which is essential for identifying potential cross-resistance mechanisms (33, 36) or stepwise progression towards high-level resistance development (23, 37, 38). Information generated from such assays has the potential to inform optimal treatment strategies for melioidosis patients in near real-time and reduce the number of cases where undiagnosed antibiotic resistance emerges during treatment (39).

RND efflux pump upregulation is common in *B. pseudomallei* strains that exhibit decreased susceptibility towards clinically relevant antibiotics. This mechanism can either by itself, or in concert with additional mutations, cause resistance towards DOX, SXT, and MEM (17, 22, 23, 24), three invaluable antibiotics in melioidosis treatment regimens. Therefore, the main goal of this study was to design, optimize and validate a triplex qPCR probe-based assay targeting the AmrAB-OprA, BpeAB-OprB, and BpeEF-OprC RND efflux pumps in *B. pseudomallei* (40). Several nucleic acid-based assays have been designed for the identification of antibiotic resistance in *B. pseudomallei* (26, 33, 34, 41,42); however, all are in singleplex format, making them unattractive for characterizing more than a handful of antibiotic resistance-conferring mutations. In contrast, probe-based PCR methods can simultaneously identify multiple targets in a single assay while being efficient, affordable, sensitive and specific.

Given the large number of mutations that can cause RND efflux pump upregulation (17, 22, 23, 28, 43), we opted for a multiplex probe-based assay format that identifies increased gene expression in favor of designing individual assays targeting each regulatory mutation, the latter of which would be both impractical (due to the large number of assays required) and prone to false negatives (due to the high likelihood of missing novel variants). In addition, detecting RNA expression ensures that efflux pump upregulation is identified without requiring a thorough understanding of the underlying regulatory networks. We first optimized the three highly specific efflux pump qPCR assays in both the singleplex and triplex formats, with all assays demonstrating high robustness and sensitivity levels, similar to the highly robust *B. pseudomallei mmsA* probe-based assay (31). Following optimization, we tested the triplex qPCR assay on 16 clinical and laboratory-generated *B. pseudomallei* mutants that had alterations in *amrR*, *bpeR*, *bpeT* or *bpeS* and which exhibited resistance towards DOX or SXT or had decreased MEM susceptibilities. We demonstrated that the triplex qPCR assay accurately quantified upregulation of efflux in all 16 regulatory mutants. Our findings confirm the value of the triplex qPCR assay for the detection of RND efflux pump upregulation.

The mechanisms conferring DOX and SXT resistance, and decreased MEM susceptibility, have recently been unraveled in *B. pseudomallei.* Importantly, all resistant strains analyzed to date have upregulated efflux pump expression (17, 22, 23, 24). Our study confirmed that upregulation of *amrB*, *bpeB* or *bpeF* was always associated with decreased MEM susceptibility. Thus, in a clinical setting, the triplex assay can be used for the accurate detection of *B. pseudomallei* isolates exhibiting decreased MEM susceptibility without the need for MIC determination. In contrast, additional mutations outside of efflux pump regulatory regions, which affect either the methyltransferase, *BPSL3085*, or the tetrahydrofolate synthesis pathway-linked enzyme pterin reductase, *folM/ptr1*, are required for the shift towards high-level DOX and SXT resistance (23, 24). The two-step nature of these mechanisms means that the triplex qPCR assay alone cannot be used to definitively determine if a strain is resistant to DOX or SXT. However, the assay can be used to rule out antibiotic resistance, as a strain with little or no efflux activity is highly likely to be sensitive to these drugs. Importantly, the ability to rapidly screen isolates and detect precursor mutations to clinically relevant DOX and SXT resistance provides a unique opportunity for health practitioners to make informed decisions regarding the suitability of current treatment, and to be aware of the risk of future resistance emergence towards these antibiotics. If increased efflux pump activity is detected in any *B. pseudomallei* strain, we strongly recommend that isolates be subjected to further MIC testing and/or the patient’s treatment regimen altered to avoid selecting for resistance emergence and subsequent treatment failure.

It has been previously shown that MEM induces the expression of AmrAB-OprA in amrR-mutated *B. pseudomallei* strains, leading to decreased MEM susceptibility (17). In the current study, the triplex qPCR assay confirmed that some, but not all, *amrR* mutants required MEM for the induction of AmrAB-OprA upregulation. In contrast, the two tested WT strains, MSHR5864 and MSHR3763, did not show an increase in the expression of *amrB* in the presence of MEM. Efflux pump expression was inducible in strains encoding in-frame, four-residue *amrR* deletions (MSHR4083 AmrR_Δ_A153-D156__ and MSHR6755 AmrR_Δ_V60-C63__), and when *amrR* was completely lost (MSHR6755 *ΔamrR);* however, induction was not required for two strains encoding either a point mutation (MSHR0292 AmrR_S174P_) or a premature stop codon (MSHR0052 AmrR_E190_^⋆^). The reason for this difference is not yet fully understood. One possibility is that RND efflux pump regulatory genes are capable of binding to substrates of the efflux systems that they regulate, and in doing so, dissociate from their promoter regions, leading to efflux pump upregulation (44, 45). This hypothesis may explain the molecular mechanism in the strains that encode the in-frame *amrR* deletions. However, this concept cannot be applied to the strain lacking *amrR*, as the regulator is not present, which would lead to efflux pump expression even in the absence of MEM. Although not examined in this study, it is possible that a second, as-yet-undiscovered regulator may also be involved in *amrAB*-*oprA* regulation. Alternately, certain mutation types or methylation patterns may reverse the function of AmrR, resulting in activation rather than repression of the operon. Further work using methods such as transcriptomics (RNA-seq) or bisulfite sequencing (methyl-seq) would be needed to confirm this hypothesis. Based on our results, we recommend that a sub-inhibitory concentration of MEM (i.e. 0.25 μg/mL) is sufficient to induce RND efflux pump upregulation in strains with decreased MEM susceptibility, without impacting RND efflux pump expression in WT strains. Although not investigated here, MEM may not be required for induction if RNA is extracted directly from clinical samples due to the preservation of native expression/methylation profiles, or even the presence of MEM *in vivo* upon sample collection.

There are some recognized limitations to our study. First, there are greater difficulties associated with handling RNA compared with DNA due to the high lability of bacterial mRNA if not protected appropriately (46). This requirement limits the application of our assay to freshly collected specimens, appropriately preserved material, or viable cultures. Second, we only assessed the use of our triplex assay on RNA extracted from subcultured isolates, and our TRIzol-based RNA extraction method is relatively slow ~4h from pelleted culture to purified and quality-controlled RNA). These factors pose a significant barrier to the rapid diagnosis of RND efflux pump upregulation. An assessment of rapid RNA extraction methods from bacterial cultures, or ideally, directly from clinical specimens (47,48), is needed to achieve same-day identification of strains with increased RND efflux expression. Finally, the complex nature of antibiotic resistance mechanisms may act as a barrier to the uptake of molecular antibiotic resistance assays in clinical laboratories due to an unfamiliarity with molecular techniques or difficulties in communicating results among healthcare professionals. The latter issue can be overcome with the availability of report templates, such as the *Mycobacterium tuberculosis* template for antibiotic resistance determination in tuberculosis patients based on WGS data, which communicates molecular results from antibiotic resistance strains to health practitioners in a clear and unambiguous manner, and without the need for specialized molecular knowledge (49).

## CONCLUSIONS

*B. pseudomallei* is an important mammalian pathogen that is endemic in most tropical and many subtropical regions. As new endemic regions continue to be unveiled, and as better recognition of this neglected pathogen filters through to the clinical setting, the number of melioidosis cases reported globally is expected to increase dramatically. Due to the high mortality rate of this disease even with antibiotic treatment, patient management should include regular screening of clinical isolates for the emerging development of antibiotic resistance to ensure optimal treatment strategies are being implemented, and to minimize the possibility of unintended treatment failure. With this goal in mind, we developed and optimized a triplex qPCR assay that simultaneously detects the expression of AmrAB-OprA, BpeAB-OprA and BpeEF-OprC in *B. pseudomallei* isolates. The triplex assay was tested on a panel of genetically characterized *B. pseudomallei* isolates of both clinical and laboratory origin, with known mutations leading to SXT or DOX resistance, or decreased MEM susceptibilities. In all *B. pseudomallei* strains that contained regulatory mutations, upregulation of at least one efflux pump was accurately detected. Rapid and facile detection of efflux upregulation is a crucial component of detecting emerging drug resistance in *B. pseudomallei*, and will aid in prompt and effective administration of individualized treatment regimens for melioidosis patients.

## MATERIALS AND METHODS

### Ethics statement

This study was approved by the Human Research Ethics Committee of the Northern Territory Department of Health and the Menzies School of Health Research (HREC 02/38).

### Melioidosis patients and corresponding *B. pseudomallei* clinical isolates

Eight Australian melioidosis cases were examined in this study. Seven cases have previously been described, including one (Pre-DPMS 89) that lacks a wild-type (WT) isogenic pair. Prior comparative genomic analysis and functional characterization showed that all non-WT strains from these patients encode RND efflux pump regulatory mutations that cause increased MICs (either resistance or decreased susceptibility) towards SXT, DOX or MEM (16, 17, 22, 23).

During this study, an adult patient, P1048, presented with acute pneumonia to Royal Darwin Hospital, Northern Territory, Australia. Initial treatment involved MEM for four weeks, followed by six weeks of CAZ and then a three-month eradication course of DOX due to SXT intolerance (neutropenia). MSHR9766 was cultured from sputum upon initial admission, and MSHR9872 was cultured from sputum during the final week of MEM treatment. P1048 reverted to being culture-negative after two months of treatment, and has since recovered from their infection. MIC testing showed that MSHR9766 was sensitive to MEM (1 μg/mL), whereas MSHR9872 had decreased susceptibility towards MEM (7 μg/mL), although both isolates were sensitive towards the other clinically relevant antibiotics (DOX, SXT and CAZ). Due to the development of an elevated MEM MIC in a latter isolate from this patient, we included this patient in our study.

The strain details for these eight cases, including regulatory mutations and MIC data, are shown in Table 1. Additionally, an *amrR* knockout of the final isolate retrieved from Patient 726 (MSHR6755 Δ*amrR*), which was previously created via allelic exchange (17), was included in the present study to determine the effect of regulator loss on efflux expression profiles.

### *B. pseudomallei* growth conditions and MIC determination

All clinical strains were grown on Luria-Bertani (LB) agar or in LB broth (Oxoid, Thebarton, SA, Australia) at 37°C for 24h unless otherwise stated. For the select agent-excluded strain Bp82 (50) and its derivatives, media were supplemented with 80 μg/mL adenine (Sigma, St. Louis, MO, USA), and for MIC testing of Bp82-derived strains, Mueller-Hinton (MH) agar or broth was supplemented with 40 μg/mL adenine. *Escherichia coli* strains DH5*α* and RHO3 were used for plasmid DNA manipulation or mobilization, respectively, and were grown according to previously published methods (51, 52). Antimicrobial susceptibility testing was performed using Etests according to the manufacturer’s instructions (bioMérieux, Baulkham Hills, NSW, Australia). Resistance cut-offs were based on the Clinical and Laboratory Standards Institute (CLSI) guidelines, as follows: CAZ S≤8, I=16 and R≥32 μg/mL; DOX: S4≤, I=8 and R≥16 μg/mL; and SXT: S≤2/38 and R≥4/76 μg/mL (53). The CLSI guidelines do not list MIC values for *B. pseudomallei* towards MEM; thus we categorized decreased MEM susceptibility as MICs ≥3 μg/mL based on prior studies (17, 54, 55). All experiments with clinical *B. pseudomallei* isolates were performed in a physical containment level 3 (biosafety level 3) facility according to local regulations, whereas experiments involving strain Bp82 and its derivatives were conducted in the physical containment level 2 laboratory, as Bp82 is excluded from select agent regulations due to an attenuated virulence phenotype conferred by a Δ*purM* mutation (www.selectagents.gov/SelectAgentsandToxinsExclusions.html).

### *B. pseudomallei* Bp82 efflux pump regulator mutants

Eight *B. pseudomallei* Bp82 strains with efflux pump regulatory mutations were included, six of which have been previously created (24): Bp82.268 Bpe_C310R_, Bp82.269 BpeT_L265R_, Bp82.270 BpeT_S280P_, Bp82.284 BpeS_P29S_, Bp82.285 BpeS_K267T_ and Bp82.280 Δ(*amrAB*-*oprA*) Δ*bpeR.* These Bp82 mutants, with the exception of Bp82.280 Δ(*amrAB*-*oprA*) Δ*bpeR*, were created via site-directed mutagenesis and knockouts to investigate the mechanisms of SXT resistance in *B. pseudomallei* using previously described methods (24). Bp82.280 Δ(*amrAB*-*oprA*)Δ*bpeR* was derived from a previously created strain (Bp82.27) as previously described (27, 56) leaving a Flp recombinase target (FRT) scar in the *bpeR* gene. MIC data for the Bp82 mutants are detailed in Table 1.

### Construction of two novel Bp82 *amrR* mutants

We have recently shown that a T520C point mutation in *amrR* (AmrR_S174P_) in an Australian clinical *B. pseudomallei* isolate plays an important role in increasing the MIC towards DOX (1 to 16 μg/mL) (23). Two laboratory-generated Bp82 *amrR* mutants (Bp82.410 Δ*amrR* and Bp82.412 AmrR_S174p_) were therefore created in this study to better understand the effect of *amrR* mutations on *amrAB*-*oprA* expression. Deletion of *amrR* was achieved using the pEXKm5-based allelic replacement system (51) (Table 2). Briefly, the US and DS region of *amrR* was amplified by PCR (using Bp82 genomic DNA as template) using primers amrR_UP_F, amrR_UP_R, amrR_DN_F and amrR_DN_R (Table 3). The PCR fragments were purified and assembled into pEXKm5 to create pEXKm5-US-DS-amrR (Table 2), using NEBuilder High-Fidelity DNA Assembly system (New England Biolabs, Ipswich, MA, USA). pEXKm5-US-DS-amrR was subsequently conjugated into Bp82 using an *E. coli* RHO3 mobilizer strain (51). Finally, merodiploids were selected for and resolved. The resultant Δ*amrR* strain was named Bp82.410. Loss of *amrR* was confirmed by PCR amplification using the primers amrR_Fullgene_F and amrR_Fullgene_R (Table 3), with the amplified size of WT *amrR* being 1,472bp compared with the *amrR* knockout at 800bp. For construction of strain Bp82.412 expressing AmrR_S174P_, the entire *amrR* gene including 400 bp upstream sequence (US) and 400 bp downstream sequence (DS) was first ligated into pGEM-T Easy (Promega, Madison, WI, USA) (Table 2) to create pGEM-T-US-DS-WT-amrR (Table 2) using primers amrR_Fullgene_F and amrR_Fullgene_R (Table 3). Site-directed mutagenesis was performed using pGEM-T-US-DS-WT-amrR together with the *amrR* mutagenic primers amrR_T520C_F and amrR_T520C_R (Table 3) to create pGEM-T-US-DS-amrRT520C (Table 2) following the manufacturer’s instructions (QuikChange II Site-Directed Mutagenesis Kit, Agilent Technologies, Santa Clara, CA, USA). The US-DS-amrR_T520C_ fragment was subcloned into the *Eco*RI site of pExKm5 creating pEXKm5-US-DS-amrR_T520C_ (Table 2). Mutagenesis was confirmed by dideoxy nucleotide sequencing of the newly introduced amrR_T520C_. The mutagenic plasmid pEXKm5-US-DS-amrR_*T*520*C*_ was then introduced into Bp82.410 Δ*amrR* as described using *E. coli* RHO3 (51), followed by allelic exchange. Merodiploids were selected for and resolved to create Bp82.412 (Bp82 AmrR_S174P_).

**TABLE 2.**
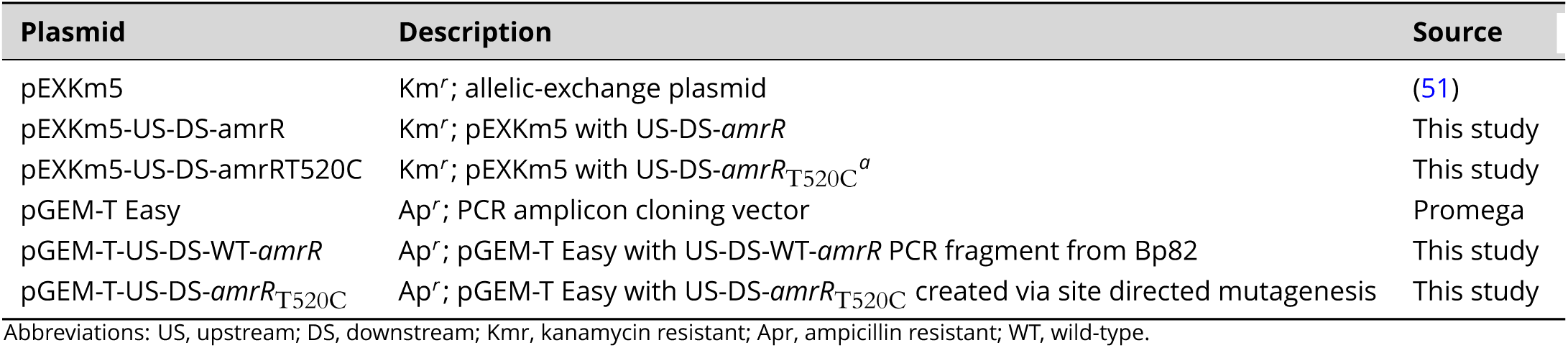
Plasmids used in the study.

| Plasmid | Description | Source |
| --- | --- | --- |
| pEXKm5 | Km<sup>r</sup>; allelic-exchange plasmid | (51) |
| pEXKm5-US-DS-amrR | Km<sup>r</sup>; pEXKm5 with US-DS-*amrR* | This study |
| pEXKm5-US-DS-amrRT520C | Km<sup>r</sup>; pEXKm5 with US-DS-*amrR*<sub>T520C</sub><sup>a</sup> | This study |
| pGEM-T Easy | Ap<sup>r</sup>; PCR amplicon cloning vector | Promega |
| pGEM-T-US-DS-WT-*amrR* | Ap<sup>r</sup>; pGEM-T Easy with US-DS-WT-*amrR* PCR fragment from Bp82 | This study |
| pGEM-T-US-DS-*amrR*<sub>T520C</sub> | Ap<sup>r</sup>; pGEM-T Easy with US-DS-*amrR*<sub>T520C</sub> created via site directed mutagenesis | This study |

Abbreviations: US, upstream; DS, downstream; Km<sup>r</sup>, kanamycin resistant; Ap<sup>r</sup>, ampicillin resistant; WT, wild-type.
<sup>a</sup>the T520C mutation corresponds to the S174P mutation present in MSHR0292.

**TABLE 3.**
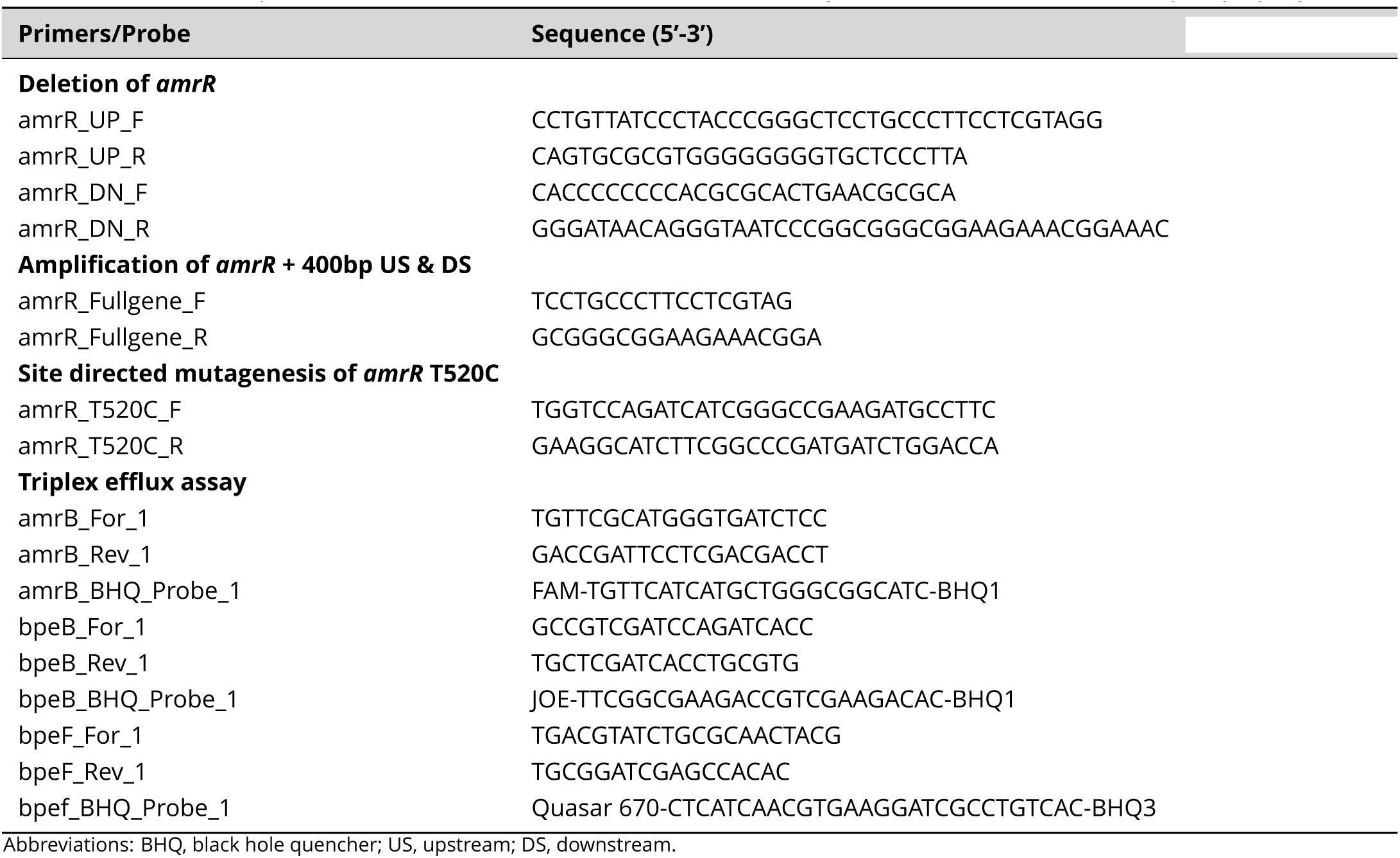
Primers and probes used for *amrR* knockout, site-directed mutagenesis and detection of efflux pump upregulation.

### Genomic analyses

We have previously used a comparative genomics approach to identify genetic variants in seven cases (16, 17, 22, 23). This same approach was also employed to identify variants between isolates obtained from a new melioidosis case, P1048, which arose during the course of our study. Reference-based assembly of the initial P1048 isolate, MSHR9766, was performed with MGAP v1.0 (https://github.com/dsarov/MGAP—Microbial-Genome-Assembler-Pipeline), using the closed genome of Australian strain MSHR1153 (57) for scaffolding. The MSHR9766 assembly was error-corrected by self-read mapping to correct for a small number of single-nucleotide polymorphism (SNP) and insertion-deletion (indel) errors. The corrected assembly was subsequently used as a reference to identify genetic variants (SNPs, indels, copy number variants) between the P1048 strains, and to rule out gene loss in the latter strain. Read mapping was carried out using BWA (58), SAMTools (59), GATK (60) and SnpEff (61), which are wrapped in the SPANDx pipeline v3.2.1 (62). Default SPANDx settings were used, with the flag for indel detection (-i) enabled. A second analysis using the Thai clinical strain *B. pseudomallei* K96243 (63) as the reference genome was performed to enable accurate variant annotation.

### RNA extractions and cDNA synthesis

To determine expression levels of the three RND efflux pumps, *B. pseudomallei* clinical strains and mutants were grown to mid-log phase (OD_600_ = 0.8 to 1) in LB broth for RNA extraction unless otherwise stated. For two *B. pseudomallei* isogenic pairs (MSHR5864 and MSHR6755; MSHR3763 and MSHR4083) and an AmrR knockout (MSHR6755 Δ*amrR*), LB was supplemented with 0.25 μg/mL MEM (Sigma-Aldrich, Castle Hill, NSW, Australia); all other strains were grown in the absence of antibiotics. For the clinical *B. pseudomallei* pairs, biological replicates of total RNA were extracted using TRIzol according to the manufacturer’s instructions (Thermo Fisher Scientific, Scoresby, VIC, Australia). RNA samples were treated with TURBO DNase (Thermo Fisher Scientific) prior to cDNA conversion using the QuantiTect Reverse Transcription Kit (Qiagen, Chadstone, VIC, Australia). Eradication of contaminating DNA prior to cDNA synthesis was confirmed with real-time PCR on the RNA extracts without the inclusion of reverse transcriptase to ensure no amplification. For the Bp82 mutants, biological triplicates of total RNA were extracted using the RNeasy Protect Bacteria kit according to the manufacturer’s instructions (Qiagen), followed by DNase treatment (Fermentas, Waltham, MA, USA) and cDNA conversion using the SuperScript III First-Strand Synthesis SuperMix (Thermo Fisher Scientific).

### Triplex assay design and quantitative PCR conditions

For the quantification of AmrAB-OprA, BpeAB-OprB and BpeEF-OprC expression, a triplex assay was designed to detect the RND transporter genes (*amrB*, *bpeB* and *bpeF*) of these efflux pumps. Primers and probes were designed using Primer Express software v3.0.1 (Applied Biosystems, Scoresby, VIC, Australia) and checked for specificity and binding efficiency using MegaBLAST. NetPrimer (http://www.premierbiosoft.com/netprimer/) was used to guide oligo design to avoid primer dimer artefacts, with a within-assay cross- and self-dimer (ΔG) cut-off of-10 or higher deemed acceptable. Each assay was first performed and optimized in singleplex format, with probes initially being labeled with the FAM dye, before converting the assays to a triplex-compatible format using three spectrally distinct dyes: 5′-FAM-amrB-BHQ-1-3′, 5′-JOE-bpeB-BHQ-1-3′ and 5′-Quasar670-bpeF-BHQ-1-3′. For *amrB* and *bpeB*, 0.35μM of each primer was used (amrB-For_1, amrB-Rev_1, bpeB-For_1, bpeB-Rev_1) with 0.2μM of *bpeF* primers, together with 0.25μM of each Black Hole Quencher probe (LGC Biosearch Technologies, Petaluma, CA, USA) and 1 XTaqMan Environmental PCR Master Mix (Applied Biosystems).

Two conserved genes were used as controls for normalizing efflux pump gene expression. For all isolates, the TaqMan MGB probe-based *mmsA* (*BPSS0619;* also known as 266152) assay, which targets methylmalonate-semialdehyde dehydrogenase (31), was used. The *mmsA* PCRs were carried out using either 384-well optical plates on the QuantStudio 6 Flex Real-Time PCR system (Thermo Fisher Scientific), or in 96-well optical plates using the CFX96 Touch Real-Time PCR detection system (Bio-Rad, Hercules, CA, USA). The limits of detection (LoD) and quantitation (LoQ) for the *mmsA* assay have previously been determined as 4 fg, or 0.5 genomic equivalents (GEs), which is the equivalent of a single PCR template (31), making this assay attractive for low-concentration target quantification. For efflux pump gene normalization in eight Bp82 mutants, a 23S rDNA assay was also used as described elsewhere (27). For 23S qPCRs, 0.2 μM of each primer was used with 2 X SYBR Select Mastermix (Thermo Fisher Scientific). The following conditions were used for thermocycling: enzyme activation for 2 min at 50°C, initial denaturation at 95°C for 10 min, followed by 45 cycles of denaturation at 95°C for 15 sec and annealing for 1 min at 60°C. The auto setting was used on each instrument when determining the threshold for each of the three assays in the triplex format.

### Triplex assay performance

The performance of the triplex qPCR assay was tested across several criteria to determine the LoQ, LoD and linearity (efficiency) following previously published methods (31, 29). Briefly, genomic DNA from *B. pseudomallei* MSHR4420 was used as template due to this sample being reasonably concentrated (286 ng/μL according to NanoDrop 2000 [Thermo Fisher Scientific] spectrophotometric analysis), of high quality and from a recent extraction. To establish the LoQ, LoD and linearity, 1:10 serial dilutions of *B. pseudomallei* MSHR4420 ranging from 40 to 4×10^‒6^ ng across eight replicates at each concentration were used as PCR template. Genomic equivalents (GEs) were calculated using an average molecular weight of 660 g/mol/bp and a 7.2 Mbp genome size.

### Triplex qPCR assay use and statistics

The triplex qPCR assay was performed on the 15 clinical strains, and eight Bp82 laboratory-generated mutants that contained regulator mutations. Average relative expression values (calculated as both times (x) change and log_2_ fold change in expression) were calculated based on biological replicates or triplicates. The relative expression data was analyzed by two-way analysis of variance (ANOVA) and Tukey’s multiple-comparison test using GraphPad Prism (GraphPad Software, Inc., La Jolla, CA). *p* values of <0.05 were considered significant.

### Availability of data and materials. WGS and accession numbers

Paired-end Illumina WGS data for the clinical strains MSHR0052, MSHR0664, MSHR0937, MSHR5651, MSHR5654, MSHR0292, MSHR0293, MSHR6522, MSHR7929, MSHR3763, MSHR4083, MSHR5864, and MSHR6755 were previously generated to ~80-90x coverage using the HiSeq2000 or HiSeq2500 platforms (Macrogen Inc., Geumcheon-gu, Seoul, Rep. of Korea). These reads have been deposited into the Sequence Read Archive database under accession numbers SRR5818275, SRR5927082, SRR2886988, SRR3381886, SRR3404570, SRR4254580, SRR4254579, SRR5949104, SRR6075126, SRR2887021, SRR2887030, SRR607512 and SRR6075122, respectively. Illumina reads for MSHR9766 and MSHR9872 were generated on the NextSeq platform as part of the current study (Macrogen Inc.), and have been deposited under accession numbers SRR6384102 and SRR6384101, respectively.

## SUPPLEMENTAL MATERIAL

**FIG S1**. Supplemental Figure S1: Range of linearity for the *Burkholderia pseudomallei* resistance-nodulation-division efflux pump triplex qPCR assay. A) *bpeB*, B) *amrB* and C) *bpeF.*

**FIG S2**. Supplemental Figure S1: Range of linearity for the *Burkholderia pseudomallei* resistance-nodulation-division efflux pump singleplex qPCR assays. A) *bpeB*, B) *amrB* and C) *bpeF.*

**TABLE S1**. Efflux pump relative expression profiles (AmrAB-OprA [*amrB*], BpeAB-OprB [*bpeB*] and BpeEF-OprC [*bpeF*]) of the Bp82 regulatory mutants. The conserved gene control (*mmsA* and/or 23S rDNA) used for each strain is indicated.

## ACKNOWLEDGMENTS

We would like to thank Linda Viberg, Ammar Aziz, Tegan Harris, Mark Mayo, Vanessa Theobald and Barbara MacHunter (Menzies School of Health Research) for laboratory assistance. This study was funded by the Australian National Health and Medical Research Council (award no. 1098337). JRW gratefully acknowledges the support provided by the Australian Federation of Graduate Women (Barbara Hale Fellowship), which made this work possible. EPP was supported by a University of the Sunshine Coast fellowship, and DSS was funded by an Advance Queensland fellowship (AQRF13016-17RD2). Work in the HPS laboratory was funded in part by National Institute of Allergy and Infectious Diseases of the National Institutes of Health (award no. U54 AI065357). The content is solely the responsibility of the authors and does not necessarily represent the official views of the National Institutes of Health. Additional funding to HPS was provided by University of Florida Preeminence start-up funds.

